# Multifaceted parental niche construction buffers microbial and competitive challenges and drives offspring dependence in burying beetles

**DOI:** 10.64898/2026.05.20.726498

**Authors:** Eric Grubmüller, Dimitri V. Meier, Melina Kreil, Lynn Schmit, Elena Scharl, Mamoru Takata, Alfons Weig, Sandra Steiger

## Abstract

Parents across diverse taxa modify the biotic or abiotic environments of their offspring. Such modifications may constitute ecological inheritance and are central to developmental niche construction, whereby organisms shape developmental conditions and selective pressures experienced by the next generation. Despite its theoretical importance, parental niche construction is often studied under simplified conditions or by focusing on single components of care, limiting our understanding of how multiple parental modifications interact in ecologically relevant contexts and shape offspring development and evolutionary trajectories. Using the burying beetle *Nicrophorus vespilloides*, we investigated how parents jointly modify chemical and microbial properties of vertebrate carcasses, a highly contested resource on which the beetle’s larvae develop. We show that under natural microbial and competitive conditions, pre-hatch parental care enhances larval survival and growth, alters cadaveric volatile emissions, and reduces carcass attractiveness to competitors. While soil type initially shapes carcass-associated microbial communities, parental care buffers these environmental effects, creating a more consistent microbiome and thereby increasing larval survival by reducing environmentally induced mortality. Larvae of the related species *Ptomascopus morio*, which lacks pre-hatch carcass preparation, showed no difference in survival between prepared and unmodified carcasses, whereas *N. vespilloides* larvae showed reduced survival on unmodified carcasses. This contrast suggests that *N. vespilloides* larvae have evolved a reliance on a parentally constructed developmental environment. Together, these findings show that parental care can constitute an integrated form of niche construction that reshapes developmental environments, enhances offspring performance, and can promote evolutionary feedbacks leading to increased offspring dependence on parental care.

## Introduction

Parental care can enhance offspring survival and growth by shaping the conditions under which offspring develop. Through behaviors such as provisioning, defense, and the regulation of physical and chemical conditions, parents influence offspring performance and the contexts in which development takes place (1, 2). When parental care modifies environmental conditions that persist over the course of offspring development, these modifications can constitute ecological inheritance, whereby offspring develop in environments shaped by parental activity (3). Niche construction theory provides a framework for understanding such environmental modification, emphasizing that organisms can actively shape biotic and abiotic conditions, thereby altering the selective regimes experienced by themselves and others (3–6). When such modifications alter the conditions under which development occurs, they constitute developmental niche construction (7). In family systems, parental care represents a specific instance of this process, when parents modify the environments in which their offspring develop, with potential consequences for their evolutionary trajectories (8, 9). Such parental niche construction may enhance resource quality, buffer environmental stressors, or reduce environmental heterogeneity experienced during development, thereby reshaping the selective pressures acting on offspring. Despite this conceptual link, the diversity and interaction of mechanisms underlying niche construction through parental care remain poorly resolved. In particular, we lack empirical studies that dissect how specific parental modifications of the offsprings’s environment alter developmental conditions through multiple pathways, how these changes reshape interactions with competitors and other organisms, and how such effects may feed back on selection acting within these modified environments.

Burying beetles provide a tractable system for examining parental niche construction in an ecologically relevant context. Reproduction occurs on small vertebrate carcasses, which serve as both the sole nutritional resource and the primary developmental environment for larvae (10, 11). These carcasses are ephemeral resources over which microbial decomposers and other scavengers compete intensely (12–15). Prior to larval hatching, parents prepare the carcass by removing fur or feathers and applying oral and anal secretions, thereby modifying both biotic and abiotic aspects of the breeding resource (10). Offspring consequently develop within an environment structured by parental activity before hatching.

Previous studies have evaluated the benefits of pre-hatch parental care by experimentally preventing parental carcass preparation and assessing the suitability of such untended carcasses for larval development under benign laboratory conditions. Despite clear evidence that parents modify carcass environments, these assays have revealed neutral (15, 16), or even negative (17, 18) effects of carcass preparation on larval performance, with positive effects reported primarily on strongly aged carcasses (15) that burying beetles rarely exploit under natural breeding conditions (15, 19, 20).

At the same time, pre-hatching care has been shown to influence specific properties of the breeding resource, including microbial community composition (21–25) and the chemical cues used by competitors for resource detection (26, 27). However, these different effects have largely been documented separately, and in different species or ecological contexts. A s a result, it remains unclear how parental modifications of the breeding resource are expressed under harsher, ecologically relevant conditions, whether they co-occur within the same developmental environment, what fitness-relevant consequences they have for larval performance, and whether the parental creation of a benign, sheltered environment influences selection acting on offspring traits.

Here, we examine parental niche construction in the burying beetle *Nicrophorus vespilloides* under ecologically relevant conditions to determine how multiple parental modifications of the breeding resource jointly shape the larval developmental environment. Combining laboratory and field experiments, we test whether the fitness consequences of pre-hatch parental care emerge when natural microbial and competitive conditions are present, and whether parental carcass preparation alters multiple, microbe-dependent properties of the breeding resource that affect larval development and interactions with competitors. Finally, using a comparative approach, we assess whether reliance on parental niche construction differs between *N. vespilloides* and a closely related species that does not engage in active pre-hatch carcass preparation, providing insight into how parental environmental modification may feed back on selection acting on offspring traits.

## Results

### Larval performance consequences of carcass preparation under naturalistic microbial conditions

We first assessed the fitness consequences of parental pre-hatch care for offspring developing on mouse cadavers placed on natural forest soil, which harbors a diverse microbial community and provides a realistic microbial challenge. We provided broods of twelve freshly hatched *N. vespilloides* larvae with cadavers that were initially fresh and had subsequently been either tended by parents or left untended for the same duration. We found that carcass preparation enhanced larval survival after 48 h (GLM, χ^2^ = 37.13, df = 1, *p* < 0.001, Fig. 1A), as well as survival to dispersal (GLM, χ^2^ = 27.25, df = 1, *p* < 0.001, Fig. 1A) and eclosion (GLM, χ^2^ =9.9, df = 1, p = 0.002; Fig. 1A). Larval mass was also higher on tended carcasses at all three time points (GLMs; 48h: χ^2^ = 5.42, df = 1, *p* = 0.02; dispersal: χ^2^ = 17.71, df = 1, *p* < 0.001; eclosion: χ^2^ = 24.08, df = 1, *p* < 0.001, Fig. 1B). Thus, parental carcass preparation strongly enhances offspring performance under natural microbial conditions.

**Figure 1:**
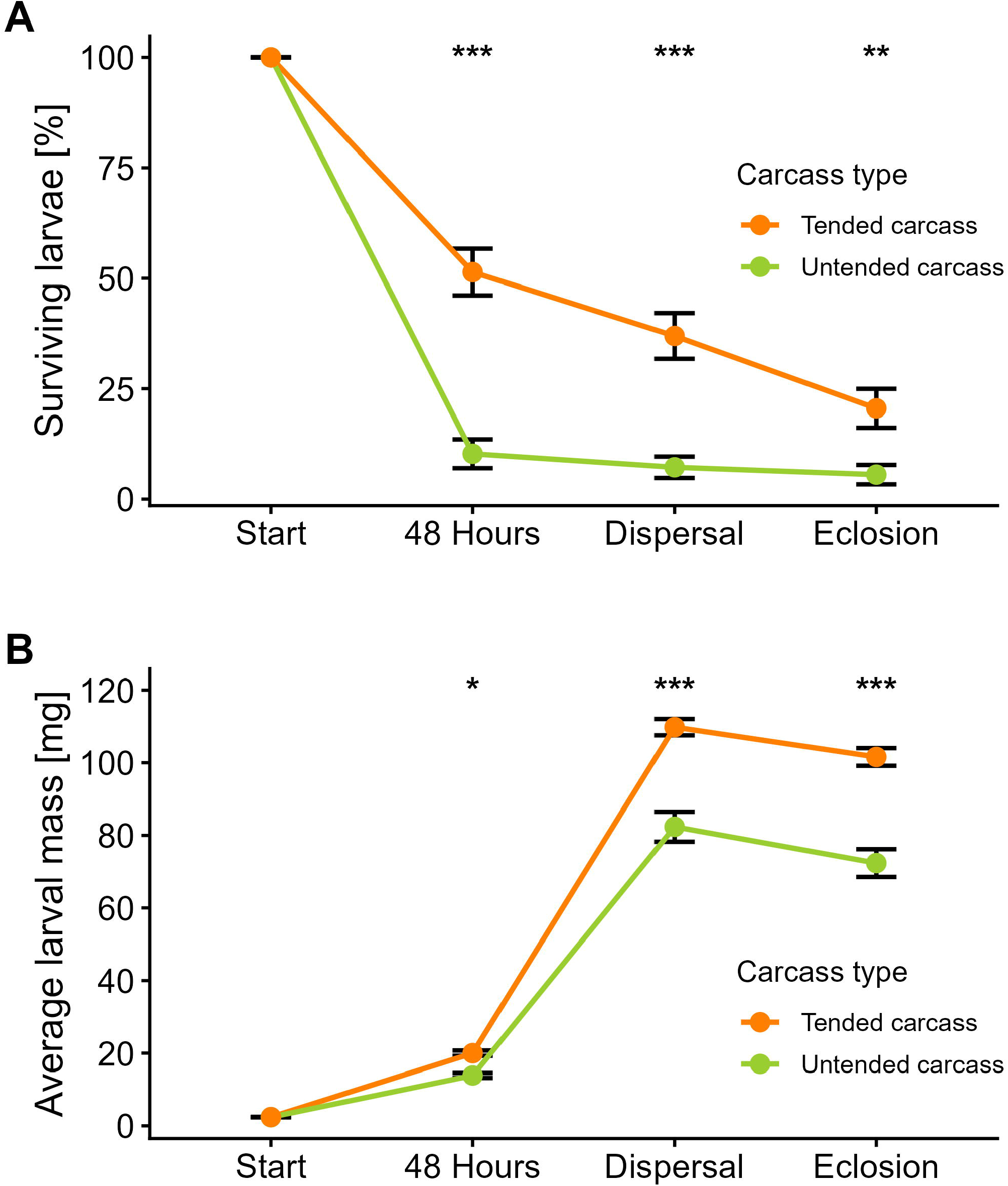
Parental pre-hatch care enhances larval survival and growth in *N. vespilloides*. Effects of parental niche construction on larval development over the entire growth period: (A) Percentage of surviving larvae relative to the original brood size; (B) Average weight of an individual larva per brood. Data are presented as mean ± SE. Asterisks indicate statistical significance (ns = not significant; * p < 0.05; ** p < 0.01; *** p < 0.001).

### Effects of carcass preparation on cadaveric VOC emissions

Since volatile organic compounds (VOCs) mediate the attraction of carrion by both vertebrate and invertebrate scavengers (28), we next investigated whether pre-hatch care altered cadaveric VOC emissions. We collected and quantified five volatiles - dimethyl disulfide (DMDS), dimethyl trisulfide (DMTS), dimethyl tetrasulfide (DMQS), S-methyl thioacetate (MeSAC) and methyl thiocyanate (MeSCN) - all sulfur-containing compounds previously shown to be affected by carcass preparation in other *Nicrophorus* species (26, 27, 29). Volatiles were sampled using a dynamic headspace method followed by GC-MS analysis. Carcass preparation affected overall VOC profiles (PERMANOVA, R^2^ = 0.15, F= 8.25, p=0.001, Fig. 2F), revealing clear differences between tended and untended carcasses. At the level of individual compounds, DMDS did not differ between tended and untended carcasses (Wilcoxon, W = 288, p = 0.82, Fig. 2A). However, tended carcasses emitted higher quantities of DMTS (Wilcoxon, W = 73.5, p<0.001, Fig. 2B) and DMQS (Wilcoxon, W = 87.5, p<0.001, Fig. 2C), but lower amounts of MeSAC (Wilcoxon, W = 420, p<0.001, Fig. 2D) and MeSCN (Wilcoxon, W = 576, p<0.001, Fig. 2E). Thus, tended carcasses emitted lower amounts of MeSCN, a known attractant (29), while increasing DMTS, previously shown to act as a deterrent in the congeneric *N. orbicollis* (29).

**Figure 2:**
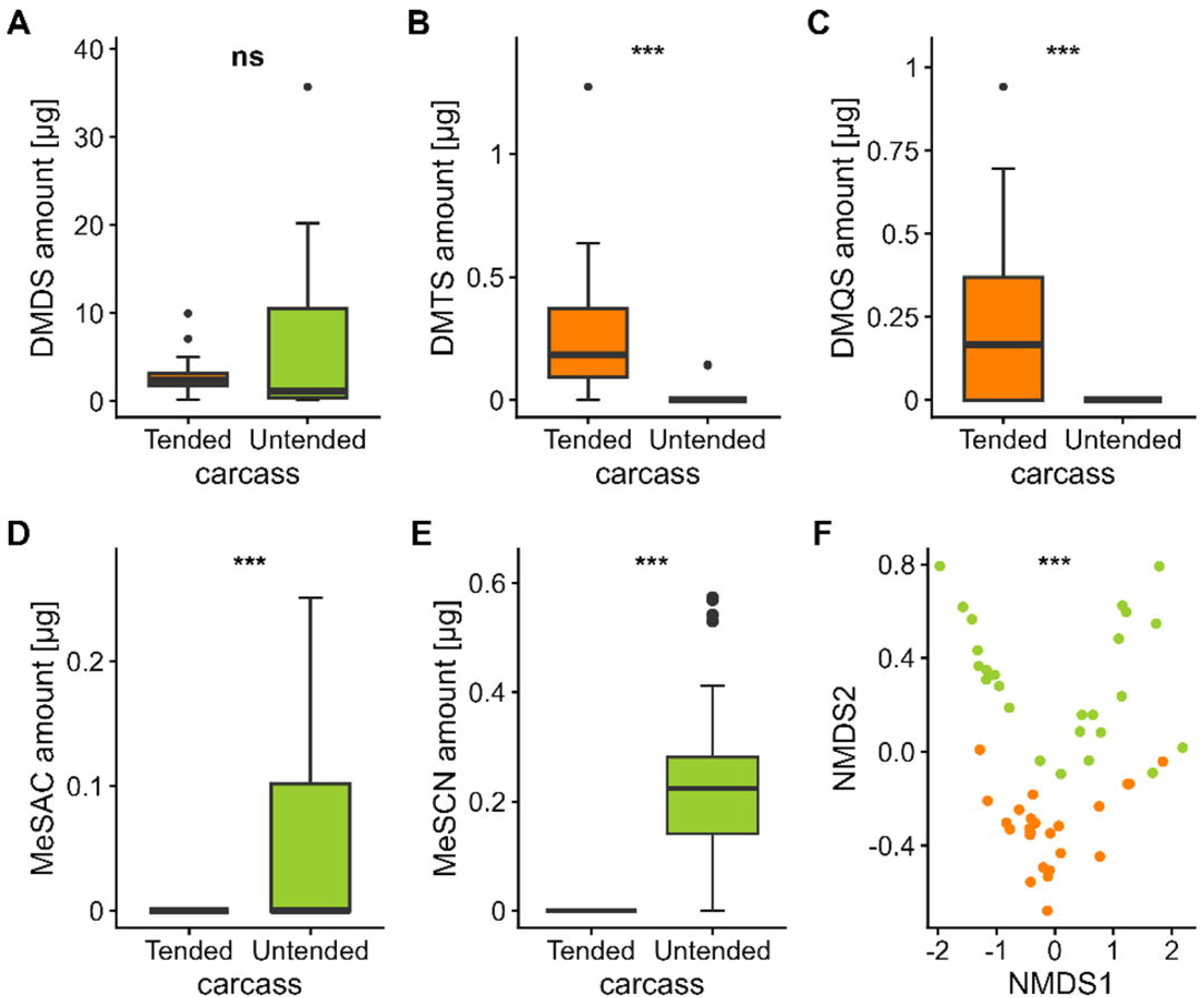
Parental pre-hatch care significantly alters the emission of cadaveric VOCs. Sulfur volatile emissions from tended and untended carcasses: (A) Dimethyl disulfide, (B) Dimethyl trisulfide, (C) Dimethyl tetrasulfide, (D) S-methyl thioacetate, and (E) Methyl thiocyanate. Boxplots represent the median and interquartile range (IQR), with whiskers extending to 1.5x IQR; individual points indicate outliers. (F) Non-metric multidimensional scaling (NMDS) ordination based on Bray-Curtis dissimilarities, illustrating the distinct chemical profiles of tended (orange) versus untended carcasses (green). Stress values: k = 2, stress = 0.035. Asterisks indicate statistical significance (ns = not significant; * p < 0.05; ** p < 0.01; *** p < 0.001).

### Effects of carcass preparation on carrion attractiveness

We then assessed the attractiveness of tended and untended carcasses to rivals in two field experiments. Because burying beetles typically bury carcasses during pre-hatch care, carcasses were presented in traps placed either above or below ground, allowing us to assess the effect of carcass preparation under both exposure conditions (see SI Appendix, Fig. S1 for trap designs). When exposed above ground, carcass preparation had no effect on the attraction of *Nicrophorus* (GLM, χ^2^ = 0.47, df = 1, p = 0.49, Fig. 3A) or silphine beetles (GLM, χ^2^ = 0.24, df = 1, p = 0.62, Fig. 3B). In contrast, dipterans were less strongly attracted to tended carcasses than to untended ones (GLM, χ^2^ = 19.30, df = 1, p < 0.001, Fig. 3C). When carcasses were presented below ground, congeneric rivals were significantly less attracted to tended than to untended carcasses (GLM, χ^2^ = 12.5, df = 1, p < 0.001, Fig. 3D). No other carrion-associated insects were recovered from the soil traps. In both experiments, neither sampling site nor sampling day (SI Appendix, Tab. S1) affected animal captures.

**Figure 3:**
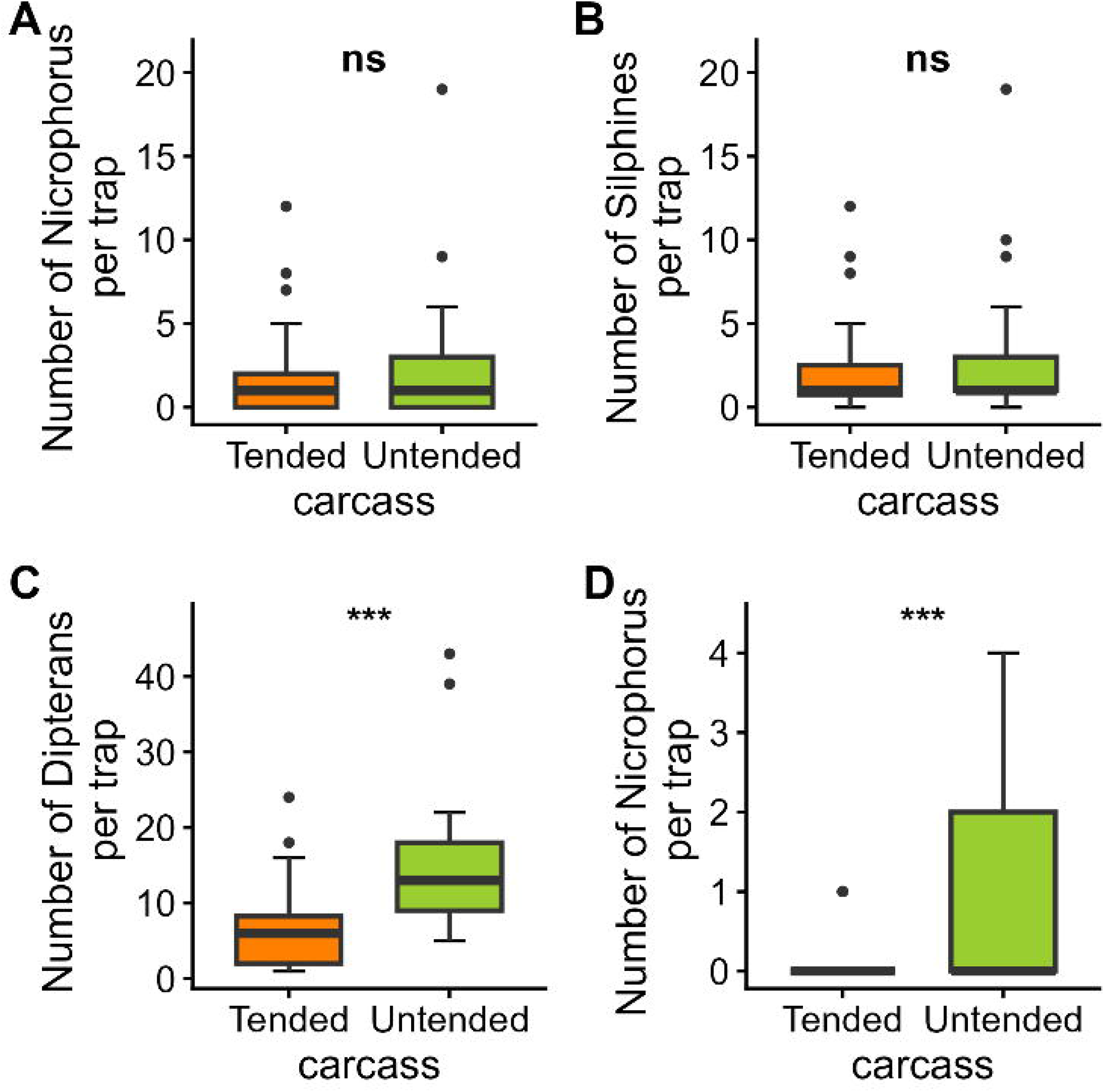
Rival avoidance is driven by chemical manipulation and carcass burial through pre-hatch care. Attractiveness (number of visitors per trap-night) of tended and untended carcasses under different exposure conditions: In experiments (A), (B), and (C), carcasses were exposed above ground; in (D), carcasses were buried below ground. Boxplots represent the median and interquartile range (IQR), with whiskers extending to 1.5x IQR; individual points indicate outliers. Asterisks indicate statistical significance (ns = not significant; * p < 0.05; ** p < 0.01; *** p < 0.001).

### Soil environment and parental care shape carrion-associated microbial communities

We next assessed how parental pre-hatch care and soil type shape carrion-associated bacterial and fungal communities using 16S rRNA and ITS amplicon sequencing. Because coconut substrate is commonly used in laboratory studies, we included it as a contrast to natural forest soil, a more complex microbial environment, to test whether carcass preparation buffers substrate-dependent differences in microbial community composition. Carcass preparation resulted in substantial changes of bacterial community compositions in both soil types. Pairwise PERMANOVA revealed that soil communities differed significantly between substrates, and carcass-associated communities varied with both carcass preparation and substrate (Fig. 4A, SI Appendix, Table S2). In particular, tended carcasses harbored distinct bacterial communities compared to untended carcasses, and bacterial communities of untended carcasses differed between substrates (Fig. 4A). In contrast, bacterial communities of tended carcasses did not differ between substrates (SI Appendix, Table S2, Fig. 4A), indicating that soil influences carcass-associated microbiota, but that carcass preparation buffers these effects. To formally test whether substrate effects depend on pre-hatch care, we then performed a second PERMANOVA restricted to carcass samples, including carcass preparation and substrate as factors. This analysis revealed a strong effect of carcass preparation (R^2^ = 0.54, F = 48.57, p < 0.001), whereas substrate alone was not significant (R^2^ = 0.03, F = 2.61, p = 0.06). Notably, a significant interaction between carcass preparation and substrate (R^2^ = 0.03, F = 2.92, p = 0.048) confirms that substrate effects depend on the presence of pre-hatch care.

**Figure 4:**
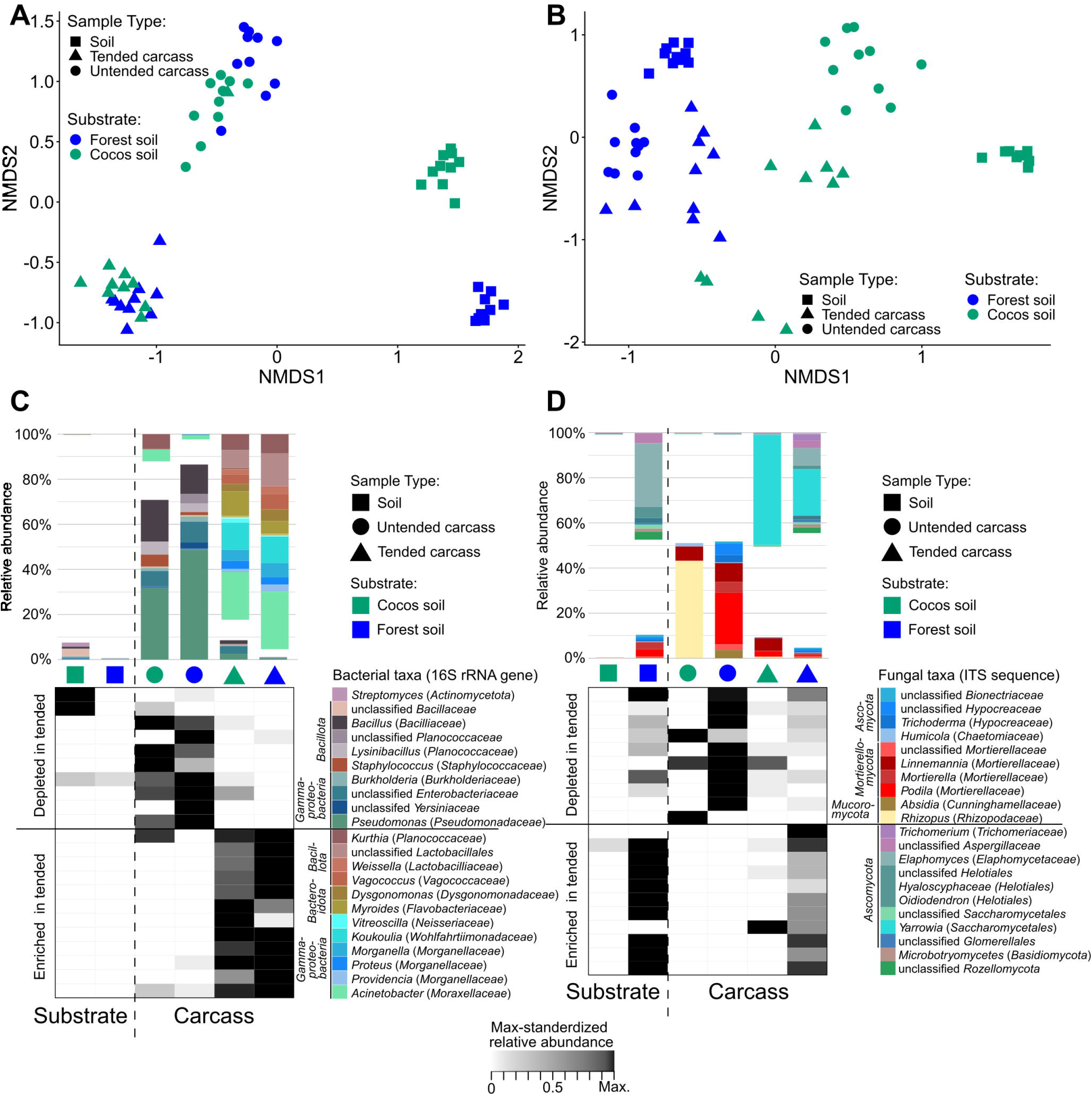
Parental niche construction overrides soil-driven succession. NMDS ordination of microbial communities based on Bray-Curtis dissimilarities for (A) bacterial and (B) fungal communities. Stress values for the ordinations are (A) k = 2, stress = 0.088; (B) k = 2, stress = 0.16. Significant differences between groups were confirmed via pairwise PERMANOVAs (SI Appendix, Tab. S2). Bar charts show the mean relative abundances (10 samples per group) of bacterial (C) and fungal (D) taxa whose marker gene sequences were significantly differentially abundant between tended and untended carcasses (DESeq2 Wald test: p ≤ 0.05, log2 fold change > 1). Taxa that were not differentially abundant or did not exceed 1% relative abundance are not shown. Heatmaps display the same taxa, with relative abundances standardized to the maximum value for each taxon to facilitate visualization of distribution patterns among low-abundance microorganisms. Relative abundance data for the entire microbial community in each individual sample are shown in Fig. S2.

Fungal communities showed overall strong differentiation among sample types. Pairwise PERMANOVA analyses across all sample types, including soil samples, revealed significant differences among all substrate and carcass groups (Fig. 4B, SI Appendix, Tab. S2), indicating that each combination of substrate and carcass preparation harbors a distinct fungal community. Hierarchical clustering and NMDS ordination (Fig. 4B and SI Appendix Fig. S2) further showed that samples grouped primarily according to substrate and secondarily by carcass preparation, suggesting that substrate represents the main axis of variation in fungal community structure. PERMANOVA analyses restricted to carcass samples identified both substrate (R^2^ = 0.22, F = 17.39, p < 0.001) and carcass preparation (R^2^ = 0.21, F = 16.11, p < 0.001) as significant predictors. Importantly, a significant interaction (R^2^ = 0.11, F = 8.36, p < 0.001) indicates that substrate effects depend on carcass preparation, as observed for bacterial communities, and are reduced in tended compared to untended carcasses. In contrast to bacteria, however, substrate-dependent differences remained detectable in tended carcasses, suggesting that carcass preparation buffers fungal communities by reducing, but not eliminating, substrate effects.

We identified the taxa that are associated with parental carcass preparation using DeSeq2. For bacterial communities, where little differences in community composition were detected between tended carcasses on forest soil and coconut substrate (Fig. 4A,SI Appendix Fig. S2A), we compared all untended versus all tended carcasses. In contrast, fungal communities were strongly substrate-dependent, as reflected in clustering patterns (Fig. 4B, SI Appendix, Fig. S2B); therefore, tended and untended carcasses were compared for each substrate separately. Tended carcasses were significantly (Log2-fold change > 1, padj ≤ 0.05) enriched in yeasts of the genus Yarrowia (Fig. 4D) and various bacterial taxa, including Bacillota (Vagococcus, Weissella, and an unclassifed family of Bacilliales), Bacteroidota (Myroides and Dysgonomonas), Gammaproteobacteria (Koukoulia, Morganella, Proteus, Providencia, and Acinetobacter) (Fig. 4C). Bacterial sequences enriched on tended carcasses, as well as Yarrowia yeast sequences, were nearly absent from soil samples and untended carcasses, indicating *N. vespilloides* exudates as their source. In contrast, many fungi enriched on tended or untended carcasses could also be found in the respective substrate, indicating that carcass preparation selects certain taxa from already present soil fungi. At the same time, tended carcasses were depleted in sequences of other Bacillota (Bacilliaceae and Planococcaceae, Staphylococcus) and Gammaproteobacteria taxa (Pseudomonas, not further classified Enterobacteriaceae and Yersiniaceae) (Fig. 4C). Fungal communities of untended carcasses were highly variable. On coconut substrate, carcasses were mostly dominated by Mucoromycota (Rhizopus) and Ascomycota (Aspergillaceae), with singular samples dominated by Mortierellomycota (Linnemannia), other Mucoromycota (Thamnidium), or Ascomycota (Pezizales) (SI Appendix Fig. S2B). Carcasses incubated on forest soil were dominated by yet distinct Mucoromycota (Mucor) and Mortierellomycota (Podila, Mortierella, and Linnemannia) sequences (SI Appendix Fig. S2B). Except for Mucor, which was present in several prepared carcasses, and the ubiquitous Aspergillaceae (SI Appendix Fig. S2B), these taxa were significantly less abundant in tended carcasses (Fig. 4D).

### Soil environment and care conditions are associated with offspring survival

To test whether the buffering effects of pre-hatch care on substrate-dependent variation in carcass microbiomes translate into fitness consequences, we measured larval survival on the same carcasses used for microbiome analyses. Larval survival until dispersal depended on an interaction between pre-hatch care and substrate (GLM, χ^2^ = 12.7, df = 1, p < 0.001). On forest soil, survival was higher on tended than on untended carcasses, whereas no effect of pre-hatch care was observed on coconut substrate (Fig. 5A, SI Appendix, Tab. S3). Although main effects of pre-hatch care (GLM, χ^2^ = 5.35, df = 1, p = 0.021) and substrate (GLM, χ^2^ = 0.00, df = 1, p = 0.99) were included in the model, the interaction best captured the observed pattern.

**Figure 5:**
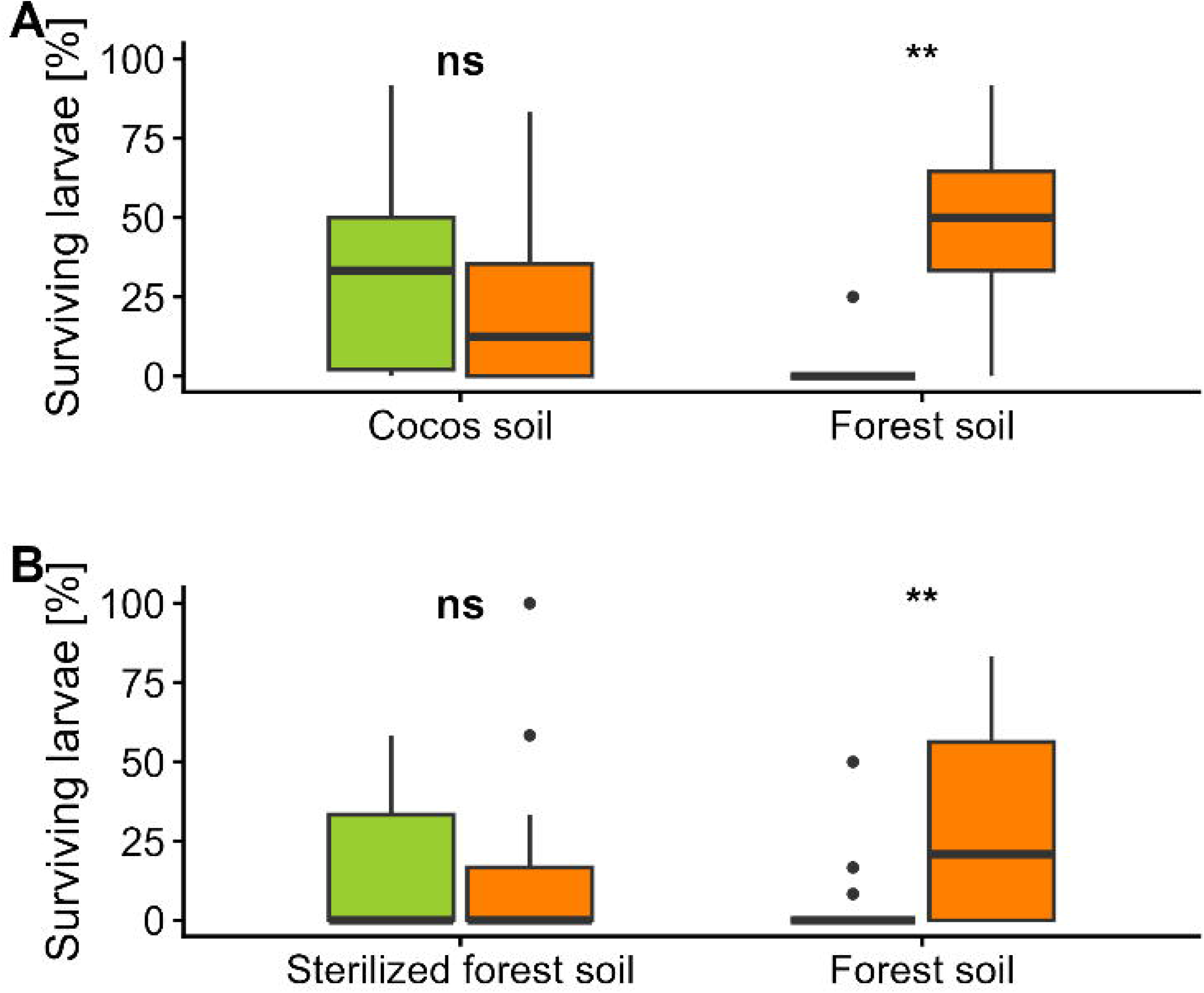
The fitness benefits of pre-hatch care are substrate-dependent and specifically critical on forest soil. Effects of parental niche construction on larval survival when raised on tended (orange) or untended (green) carcasses in (A) either cocos substrate or forest soil or (B) untreated or sterilized forest soil. Boxplots represent the median and interquartile range (IQR), with whiskers extending to 1.5x IQR; individual points indicate outliers. Asterisks indicate statistical significance (ns = not significant; * p < 0.05; ** p < 0.01; *** p < 0.001).

### Specific microbial taxa are associated with high-mortality conditions

To identify candidate microbial drivers of reduced offspring performance, we focused on taxa significantly enriched in untended carcasses on forest soil, where high larval mortality was observed, compared to both tended carcasses on forest soil and untended carcasses on coconut substrate, where mortality was lower. Among bacteria, only five low-abundance taxa were specifically enriched in untended carcasses on forest soil. Of these, only unclassified Planococcaceae (Bacilliales) exceeded 1% mean relative abundance (4.3% ± 3.8%). Unclassified Yersiniaceae (Enterobacterales) also showed their highest relative abundance in untended carcasses on forest soil (2.6% ± 1.8%; Fig. 4C), but their increase relative to untended carcasses on coconut substrate was not significant (log2 fold change = 3.5, p = 0.09). Among fungi, 15 taxa were significantly more abundant in untended carcasses on forest soil than in both tended carcasses on forest soil and untended carcasses on coconut substrate. Six exceeded 1% relative abundance, with Podila (Mortierellomycota) dominating (23% ± 13%). Other enriched taxa included Mortierella (4.8% ± 3%), Absidia (2.8% ± 2%), and unclassified Hypocreaceae (2.7% ± 0.9%) (Fig. 4D). These results suggest that specific soil fungi play a major role in reduced survival success of the larvae.

### Soil sterilization confirms microbial mediation of offspring mortality

To directly test whether the mortality difference between soil types was driven specifically by soil-associated microbes rather than other properties of the substrate, we compared larval survival on tended and untended carcasses placed on untreated versus heat-sterilized forest soil. Larval survival depended on the interaction between pre-hatch care and soil sterilization (GLM, χ^2^ = 7.75, df = 1, p = 0.005). On untreated forest soil, larvae exhibited higher survival on tended than on untended carcasses (Fig. 5B,SI Appendix, Tab. S3), whereas this difference disappeared when the soil was sterilized (Fig. 5b, SI Appendix, Tab. S3). Although main effects of pre-hatch care (GLM, χ^2^ = 7.27, df = 1, p = 0.007) and substrate type (GLM, χ^2^ = 0.38, df = 1, p = 0.54) were included in the model, the interaction best captured the observed pattern. Taken together, these results show that carcass preparation enhances offspring survival in a substrate-dependent manner and that this effect is mediated by soil-associated microbes, indicating that pre-hatch care buffers the negative effects of microbially challenging environments.

### Species–specific effects of pre-hatch care on offspring performance

Lastly, we examined how parental pre-hatch care may feed back on selection acting on offspring traits, by comparing the performance of offspring of *N. vespilloides* and its close relative *Ptomascopus morio*, a species that lacks active pre-hatch carcass preparation. We placed twelve larvae of either species on carcasses that had been either tended by *N. vespilloides* parents or left untended on forest soil and monitored survival and mass at 48 h and at dispersal. Species effects were consistently strong, with *P. morio* showing higher survival than *N. vespilloides* at both 48 h and dispersal (GLMs, 48 h: χ^2^ = 288.31, df = 1, *p* < 0.001; dispersal: χ^2^ = 299.71, df = 1, *p* < 0.001; Fig. 6A, SI Appendix, Tab. S4). Carcass preparation also significantly affected survival (GLMs, 48 h: χ^2^ = 11.17, df = 1, *p* < 0.001; dispersal: χ^2^ = 11.91, df = 1, *p* < 0.001, Fig. 6A,SI Appendix Tab. S4). However, this effect of pre-hatch care was driven by *N. vespilloides*: while survival of *P. morio* was unaffected by carcass preparation, *N. vespilloides* showed markedly reduced survival on untended carcasses (Fig. 6A, SI Appendix Tab. S5). The same pattern was observed across development, with reduced survival of *N. vespilloides* on untended carcasses but no effect of carcass preparation in *P. morio* (Fig. 6A, Appendix Tab. S5), although the interaction between species and pre-hatch care reached statistical significance only at dispersal (GLM, χ^2^ = 4.17, df = 1, *p* = 0.041, SI Appendix Tab. S4). Larval mass did not show the same species-specific response to pre-hatch care as survival. Both species and pre-hatch care significantly influenced mass at 48 h (GLM, Species: χ^2^ = 102.49, df = 1, *p* < 0.001; pre-hatch care: χ^2^ = 5.27, df = 1, *p* = 0.022; Fig. 6B, SI Appendix, Tab. S4) and at dispersal (GLM, Species: χ^2^ = 151.41, df = 1, *p* < 0.001; pre-hatch care: χ^2^ = 13.75, df = 1, *p* < 0.001; Fig. 6B, SI Appendix, Tab. S4). At 48 h, both species showed only a tendency to be heavier on tended carcasses, but by dispersal larvae of both species were consistently heavier when reared on tended carcasses (Fig. 6B, SI Appendix, Tab. S5). No significant interactions between species and pre-hatch care were detected for mass at any stage (SI Appendix, Tab. S4).

**Figure 6:**
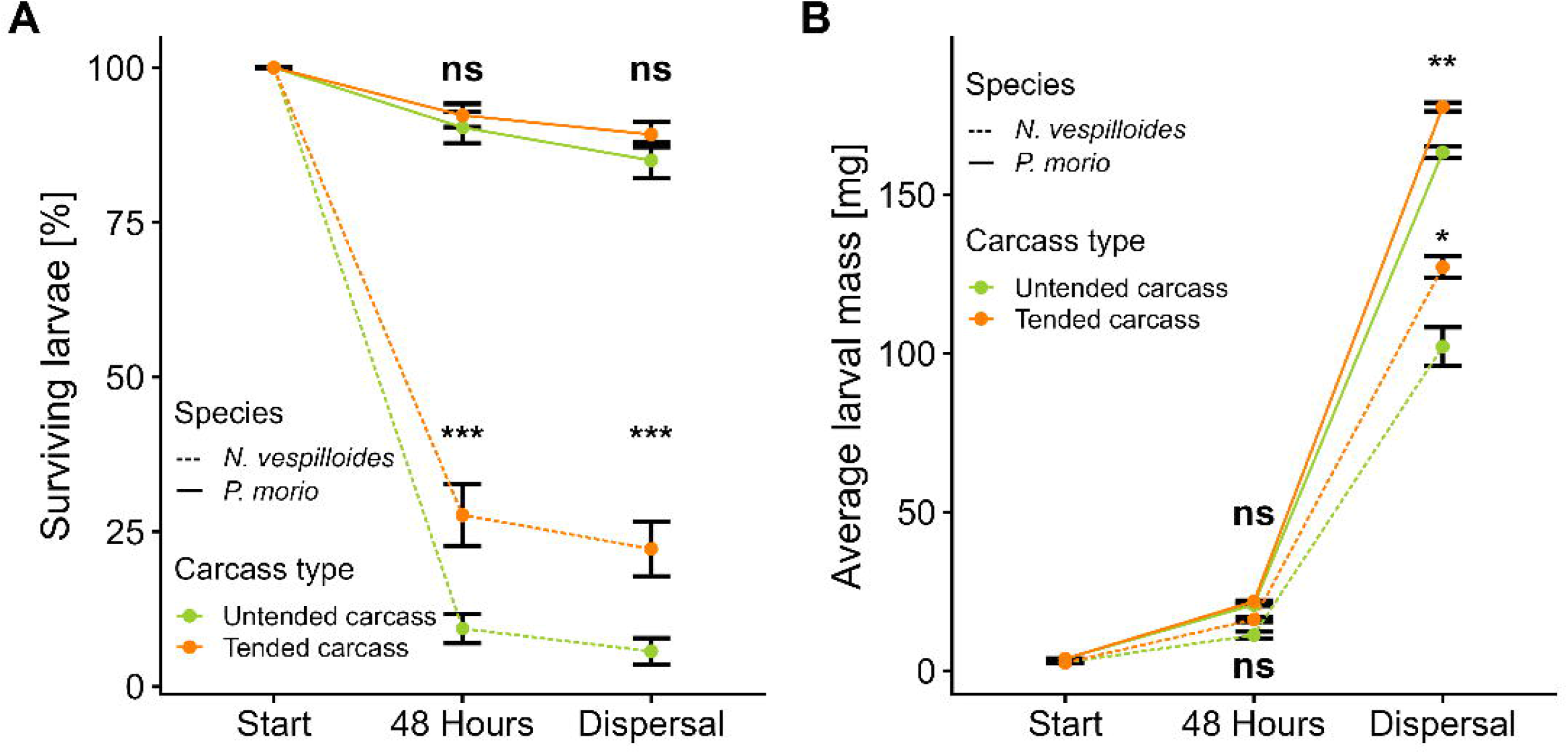
*P. morio* does not rely on carcass preparation for survival but benefits from parental niche construction through increased growth. Effects of parental pre-hatch care on (A) larval survival and (B) larval growth of *N. vespilloides*. The data are presented as mean ± SE. Asterisks indicate statistical significance between carcass types within species (ns = not significant; * p < 0.05; ** p < 0.01; *** p < 0.001).

## Discussion

By examining reproduction within a more natural microbial and competitive context, our results show that pre-hatch care in *N. vespilloides* constitutes an integrated form of parental niche construction, involving multiple, interacting modifications of the breeding resource. Under these conditions, tended carcasses supported substantially higher larval survival and growth, revealing fitness benefits that are not apparent under simplified laboratory settings. These advantages are accompanied by coordinated parental modifications of the breeding resource. Parents alter microbially mediated volatile emissions, reducing the attractiveness of the carcass to competitors, and restructure microbial communities in ways that suppress harmful organisms and buffer environmentally induced variation. Together, these processes reshape carrion into a buffered and less contested developmental environment. Notably, *N. vespilloides* larvae performed poorly on untended carcasses, whereas a closely related species lacking pre-hatch carcass preparation did not, suggesting an evolved reliance on a parentally constructed developmental environment. This pattern is consistent with the idea that niche construction can generate evolutionary feedback promoting offspring dependence on parentally modified environments.

The fitness consequences of pre-hatch care became apparent only when larvae developed on forest soil, a substrate that introduces a diverse and ecologically relevant microbial community. Under these conditions, carcass preparation substantially increased larval survival and mass gain, whereas no comparable effect was observed on coconut substrate within our experiment, indicating that the benefits of carcass preparation depend on the microbial context and helping to explain why earlier studies did not detect positive effects of carcass preparation under simplified laboratory conditions (16–18). As resource quantity was not limiting in our experiment (30), reduced performance on unprepared carcasses is unlikely to reflect resource depletion alone but instead points to costs arising from microbial activity that degrades carcass quality. Soil microbial communities are known to strongly influence the rate and progression of carrion decomposition (31), potentially including the production of toxins or other microbial byproducts associated with competition among decomposers (13). Our findings suggest that the benefits of pre-hatch care emerge through interactions with complex microbial communities that are absent or reduced under standard laboratory conditions.

Pre-hatch care also altered cadaveric VOC emissions, suggesting a mechanism by which parents modify how the resource is perceived by competitors. While DMDS release remained unchanged, parental preparation markedly increased DMTS and DMQS and nearly eliminated MeSAC and MeSCN. Previous work shows that DMDS and MeSAC dominate early decomposition and then decline, whereas DMTS and DMQS characterize later stages (32). MeSCN is a strong attractant for burying beetles and may signal high resource quality (29), whereas DMTS has been associated with reduced attraction in at least one burying beetle species (26). Together, these shifts may indicate that parents alter the apparent decay stage of the carcass, making it resemble a more advanced stage of decomposition. This likely reduces competition, as burying beetles typically prefer fresher carcasses when selecting breeding resources (15, 19).

Field assays support this interpretation. Below ground, tended carcasses attracted fewer congeneric rivals, thereby reducing competition and the associated risk of carcass takeover and brood loss through infanticide (33, 34). Above ground, tended carcasses attracted fewer Diptera but similar numbers of carrion beetles, suggesting that reduced competition from carrion beetles depends on the combination of burial and carcass preparation. Flies also showed reduced attraction to tended carcasses, despite DMTS generally being a potent attractant for many fly species (35–38). This might reflect that VOCs signal a later-stage or beetle-prepared resource that is less attractive to many fly species, either because they prefer early stages or avoid occupied carcasses.

The changes in VOC emissions are likely linked to underlying shifts in the microbial community of the carcass. Our results show that parental care suppresses components of the carcass-associated microbiota while introducing characteristic microbial taxa from parental guts and exudates and selecting for specific taxa of soil derived fungi. Together these processes establish a microbiome resembling that is associated with *N. vespilloides*, including *Yarrowia* and dominant bacterial genera such as *Vagococcus, Weissella* (both *Lactobacilliales*), *Myroides* (*Flavobacteriales*) and *Dysgonomonas* (*Bacteroidales*) (21, 23–25, 39). Several of these microbes are associated with antimicrobial activity or modification of the carrion resource (40), with *Yarrowia* in particular being a well-established symbiont of burying beetles that has been implicated in carrion predigestion, thereby increasing resource accessibility (23, 39, 41). Some of these taxa are also known to produce sulfur-containing volatiles such as DMTS (42, 43), indicating a direct microbial contribution to the altered VOC profile. Environmental microbial communities are known to strongly influence carrion decomposition and carcass-associated microbial communities (31), but parental preparation reduced substrate-dependent differences in bacterial communities and, to a lesser extent, in fungal communities, indicating convergence across environments. This microbial buffering was also reflected in larval performance: substrate-dependent differences in survival on untended carcasses disappeared when carcasses were tended. This parallel suggests that parental care stabilizes the microbial environment and its consequences for offspring. Supporting this interpretation, untended carcasses on forest soil were enriched with taxa associated with detrimental or pathogenic effects, including *Mucor, Podila*, and members of the Yersiniaceae (44, 45). Overall, these findings indicate that parental care buffers offspring development against substrate-dependent microbial variation.

In *P. morio*, which lacks carcass preparation (46, 47), larval survival was unaffected by carcass preparation, indicating that unmodified resources remain suitable, as is typical for many carrion-feeding insects that readily exploit decomposing resources (37, 48). In contrast, *N. vespilloides* larvae performed more poorly on unprepared carcasses, suggesting a reliance on a parentally constructed developmental environment. This pattern is consistent with evolutionary feedbacks under niche construction that relax selection on larval self-sufficiency, potentially reducing investment in traits required to cope with unmodified carcasses, such as microbial defense (6). At the same time, *P. morio* larvae achieved higher body mass on tended than untended carcasses, indicating that parental niche construction confers direct performance benefits, likely by reducing microbial competition for the resource and limiting degradation of carrion quality. Modification of the resource thus creates a developmental niche that enhances larval growth more generally and may therefore have favored the evolution of carcass preparation. Together, these results suggest an evolutionary trajectory in which parental niche construction initially provides facultative benefits but subsequently promotes offspring dependence on the modified environment.

Environmental modification by parents, including nest building and resource preparation, is widespread across taxa (49–52), yet its often multicomponent nature and joint effects on developmental conditions, offspring fitness, and evolutionary dynamics remain to be investigated in an integrated, experimental framework. Here, we show that parental pre-hatch care in *N. vespilloides* operates through multiple niche-constructing processes that reshape and buffer the developmental environment. In this system, larvae develop on small, ephemeral carcasses that can be monopolized and maintained by parents, such that investment in preserving resource quality and excluding competitors is favored, promoting the evolution of elaborate niche construction. Some modifications, such as carcass burial, carcass shaping and the introduction of microbial symbionts, contribute to inceptive niche construction (5, 53) by establishing a novel developmental environment relative to unmodified carrion. At the same time, these parentally induced changes also function in a counteractive manner (“counteractive niche construction” (5, 53), as they suppress detrimental organisms and reduce competition, buffering parents and offspring against pre-existing threats rather than creating new selective environments. Together, our findings demonstrate how parental behavior can both construct and stabilize the developmental niche through multicomponent effects, enhances offspring performance, and can generate evolutionary feedback that promotes increased offspring dependence on parentally modified environments.

## Methods

### Experimental animals and carcasses

*Nicrophorus vespilloides* and *Ptomascopus morio* beetles derived from lab-bred populations reared in climate chambers (20°C, 16:8 h photoperiod). Using mouse carcasses (10 ± 2 g), we established two carcass types: tended carcasses, processed by a pair of burying beetles, and untended carcasses, incubated in soil to mimic the decay status of an untended carcass at larval emergence. Carcasses were placed either on the humus-rich upper layer of forest soil, collected in a mixed forest near Bayreuth, Germany or on coconut substrate (Tropic Shop, Nordhorn, Germany), commonly used in our lab. (SI Appendix, SI Methods)

### Larval Performance on Forest Soil

We assessed the effects of parental pre-hatch care on offspring performance by offering twelve freshly hatched *N. vespilloides* larvae to mouse cadavers placed on forest soil that had either been tended by *N. vespilloides* parents (N = 30) or left untended (N = 30). We quantified larval performance by weighing larvae before adding them to the carcass and by counting and weighing all surviving larvae after 48 hours, at dispersal and at adult emergence. Larvae were considered dispersed when all individuals in a sample had left the carcass to pupate.

### Cadaveric VOCs

We analyzed volatile organic compounds (VOCs) emitted from tended (N = 24) and untended (N = 25) carcasses placed on forest soil using dynamic headspace sampling. Each carcass was enclosed in an oven bag, and filtered air was pumped through at a constant flow for one hour at room temperature. Volatiles were trapped on activated charcoal filters and subsequently extracted with dichloromethane containing methyl undecanoate (20 ng/ µl) as an internal standard. Extracts were injected splitless into a GC-MS system equipped with a polar capillary column. Compounds were identified by comparison with mass spectral libraries and retention times of reference standards. (SI Appendix, SI Methods).

### Field experiment

We examined the attractiveness of tended and untended carcasses to insect rivals in two field experiments (N = 27 carcasses per treatment per experiment). Carcasses were exposed either above or below ground. For above-ground exposure, carcasses were placed in perforated plastic cups mounted above pitfall traps. For below-ground exposure, carcasses were placed in soil-filled flower pots and covered with 4 cm of soil; pots were inserted into the ground so that their rims were level with the surface. For both experiments, we established 18 traps in a mixed forest in Bavaria, Germany. We rotated treatments daily to prevent locational bias. All captured burying beetles (*Nicrophorus* spp.), other silphine beetles (Silphinae), and dipteran insects were collected and counted (SI Appendix, SI Methods, Fig. S1).

### Characterization of the Carcass Microbiome

We characterized microbial communities associated with carcasses, forest soil, and coconut substrate using 16S rRNA and ITS2 amplicon sequencing. Carcasses exposed to the two substrate types were sampled after 72 h, either following pre-hatch care by *N. vespilloides* or after untended incubation (N = 10 per per carcass and soil type). Microbial material was collected by swabbing feeding cavities of tended carcasses, or the equivalent skin region of untended carcasses, and immediately preserved in RNAlater stabilization medium (Sigma-Aldrich, Steinheim, Germany). Substrate samples were collected at setup (N = 10 per substrate type). DNA was isolated using the DNAeasy Power Soil kit (Qiagen, Hilden, Germany). The bacterial V4 region of the 16S rRNA gene was amplified using primers 515F-Y and 806RB, and the fungal ITS2 region using primers gITS7 and ITS4. Amplicons were purified, indexed in a secondary PCR, pooled at equimolar concentrations, size-selected, and sequenced in single-end mode on an iSeq-100 Illumina platform. The 16S rRNA and ITS amplicon sequence data were analyzed using DADA2 (54) in QIIME2 (55) Taxonomic classification of ASVs was obtained using naïve-bayes trained classifiers based on the SILVA 138.2 (for 16S amplicons (56)), and UNITE 10 (for ITS2 amplicons (57)), respectively (SI Appendix, SI Methods, SI Tab. 6).

### Larval survival across care conditions and substrates

We tested how pre-hatch care and substrate affect larval survival in the absence of post-hatch care by offering twelve larvae to the same carcasses used for microbiome analyses. The experiment comprised tended and untended carcasses placed on forest soil and coconut substrate (tended-coconut: N = 12; tended-forest: N = 12; untended-coconut: N = 10; untended-forest: N = 10). The increased sample size in the tended treatments reflects two additional carcasses that were prepared as backups for potential brood failure and used only for larval fitness measurements. Additionally, to assess the effect of substrate sterilization, we compared tended and untended carcasses placed on either untreated forest soil or sterilized forest soil (tended-sterilized: N = 21; tended-forest: N = 22; untended-sterilized: N = 21; untended-forest: N = 23) (See SI Appendix, SI Methods for soil sterilization). In both experiments, we counted the number of surviving larvae at dispersal.

### Species–specific effects of pre-hatch care on offspring performance

We investigated how pre-hatch care influences larval survival and growth in *P. morio* and *N. vespilloides* in the absence of post-hatch care. To do this, we offered twelve larvae to tended (*P. morio*: N = 27, *N. vespilloides:* N = 31) or untended (both species:-N = 25) carcasses placed on forest soil. Because oviposition and egg development take longer in *P. morio*, we established pairs of this species two days earlier than *N. vespilloides*. After oviposition and before larval hatching (approximately 96 hours in *P. morio* and 48 hours in *N. vespilloides*) we transferred carcasses and beetles into fresh boxes filled with forest soil. On the following day, we monitored for hatching larvae. Larvae were pooled by species and assigned to carcasses that had either been tended by *N. vespilloides* or incubated in soil. We assessed larval performance by weighing larvae before placement on the carcass and then recording survival and individual body mass at two stages: 48 hours after introduction, and at dispersal.

### Statistical analysis

All statistical analyses were performed in R (version 4.4.2). Model fits were validated using *DHARMa* (58), no major violations were detected (SI Appendix, SI Methods).

## Data availability

Raw amplicon sequence reads have been deposited in the European Nucleotide Archive under project number PRJEB112826. All processed data tables and R scripts used for the analyses are available on **XX** at DOI: **XX**.

## Acknowledgements

We thank A. Kirpal, D. Lauterbach and J. Adler for laboratory assistance and maintenance of beetle populations, J. Stökl for his assistance in the chemical analysis, J. Kramer for his helpful comments on the manuscript and the entire team of The Evolutionary Animal Ecology Department, University of Bayreuth, for their support. This work was funded by the German Research Foundation (DFG) to S.S. (STE 1874/6-3; Project number: 277139873).

## References

1. T. H. Clutton-Brock, The Evolution of Parental Care (Princeton University Press, 1991).

2. N. J. Royle, P. T. Smiseth, M. Kölliker, Eds., The evolution of parental care (Oxford University Press, 2012).

3. J. Odling-Smee, D. H. Erwin, E. P. Palkovacs, M. W. Feldman, K. N. Laland, Niche Construction Theory: A Practical Guide for Ecologists. Q. Rev. Biol. 88, 3–28 (2013).

4. K. Laland, B. Matthews, M. W. Feldman, An introduction to niche construction theory. Evol. Ecol. 30, 191–202 (2016).

5. K. Laland, J. F. Odling-Smee, J. Endler, Niche construction, sources of selection and trait coevolution. Interface Focus 7, 20160147 (2017).

6. B. Matthews, et al., Under niche construction: an operational bridge between ecology, evolution, and ecosystem science. Ecol. Monogr. 84, 245–263 (2014).

7. D. B. Schwab, S. Casasa, A. P. Moczek, Evidence of developmental niche construction in dung beetles: effects on growth, scaling and reproductive success. Ecol. Lett. 20, 1353–1363 (2017).

8. E. G. Flynn, K. N. Laland, R. L. Kendal, J. R. Kendal, Target Article with Commentaries: Developmental niche construction. Dev. Sci. 16, 296–313 (2013).

9. T. Uller, “Parental effects in development and evolution” in The Evolution of Parental Care, N. J. Royle, Smiseth Per T., M. Kölliker, Eds. (Oxford University Press, 2012), pp. 247–266.

10. A. L. Potticary, et al., Revisiting the ecology and evolution of burying beetle behavior (Staphylinidae: Silphinae). Ecol. Evol. 14, e70175 (2024).

11. E. Pukowski, Ökologische untersuchungen an necrophorus f. Z. Für Morphol. Ökol. Tiere 27, 518–586 (1933).

12. T. L. DeVault, O. E. Rhodes, Jr., J. A. Shivik, Scavenging by vertebrates: behavioral, ecological, and evolutionary perspectives on an important energy transfer pathway in terrestrial ecosystems. Oikos 102, 225–234 (2003).

13. D. H. Janzen, Why Fruits Rot, Seeds Mold, and Meat Spoils. Am. Nat. 111, 691–713 (1977).

14. M. Körner, S. Steiger, S. P. Shukla, Microbial management as a driver of parental care and family aggregations in carrion feeding insects. Front. Ecol. Evol. 11, 1252876 (2023).

15. D. E. Rozen, D. J. P. Engelmoer, P. T. Smiseth, Antimicrobial strategies in burying beetles breeding on carrion. Proc. Natl. Acad. Sci. 105, 17890–17895 (2008).

16. A.-K. Eggert, M. Reinking, J. K. Müller, Parental care improves offspring survival and growth in burying beetles. Anim. Behav. 55, 97–107 (1998).

17. A. Capodeanu-Nägler, et al., From facultative to obligatory parental care: Interspecific variation in offspring dependency on post-hatching care in burying beetles. Sci. Rep. 6, 29323 (2016).

18. P. Huber, S. Bartl, J. Schneider, S. Steiger, Better Together: Offspring Benefit from Siblings in Both the Absence and the Presence of Parents. Am. Nat. 206, 285–297 (2025).

19. P. J. Delclos, T. L. Bouldin, J. K. Tomberlin, Olfactory Choice for Decomposition Stage in the Burying Beetle Nicrophorus vespilloides: Preference or Aversion? Insects 12, 11 (2020).

20. S. T. Trumbo, Fate of mouse carcasses in a N orthern W oodland. Ecol. Entomol. 41, 737–740 (2016).

21. A. Duarte, M. Welch, C. Swannack, J. Wagner, R. M. Kilner, Strategies for managing rival bacterial communities: Lessons from burying beetles. J. Anim. Ecol. 87, 414–427 (2018).

22. C. J. Miller, S. T. Bates, L. M. Gielda, J. C. Creighton, Examining transmission of gut bacteria to preserved carcass via anal secretions in Nicrophorus defodiens. PLOS ONE 14, e0225711 (2019).

23. S. P. Shukla, et al., Microbiome-assisted carrion preservation aids larval development in a burying beetle. Proc. Natl. Acad. Sci. 115, 11274–11279 (2018).

24. S. P. Shukla, H. Vogel, D. G. Heckel, A. Vilcinskas, M. Kaltenpoth, Burying beetles regulate the microbiome of carcasses and use it to transmit a core microbiota to their offspring. Mol. Ecol. 27, 1980–1991 (2018).

25. H. Vogel, et al., The digestive and defensive basis of carcass utilization by the burying beetle and its microbiota. Nat. Commun. 8, 15186 (2017).

26. S. T. Trumbo, P. K. B. Philbrick, J. Stökl, S. Steiger, Burying Beetle Parents Adaptively Manipulate Information Broadcast from a Microbial Community. Am. Nat. 197, 366–378 (2021).

27. S. T. Trumbo, E. Grubmüller, S. Steiger, Species-specific manipulation of microbially derived volatiles by burying beetle parents. Ecol. Entomol. een.70085 (2026). 10.1111/een.70085.

28. J. A. Cammack, M. L. Pimsler, T. L. Crippen, J. K. Tomberlin, “Chemical Ecology of Vertebrate Carrion” in Carrion Ecology, Evolution, and Their Applications, (CRC Press, 2015), pp. 187–203.

29. S. T. Trumbo, S. Steiger, Finding a fresh carcass: bacterially derived volatiles and burying beetle search success. Chemoecology 30, 287–296 (2020).

30. J. Bartlett, C. M. Ashworth, Brood size and fitness in Nicrophorus vespilloides (Coleoptera: Silphidae). Behav. Ecol. Sociobiol. 22, 429–434 (1988).

31. C. L. Lauber, et al., Vertebrate Decomposition Is Accelerated by Soil Microbes. Appl. Environ. Microbiol. 80, 4920–4929 (2014).

32. B. Kalinová, H. Podskalská, J. Růžička, Irresistible bouquet of death—how are burying beetles (Coleoptera: Silphidae: Nicrophorus) attracted by carcasses. Naturwissenschaften 96, 889–899 (2009).

33. S. T. Trumbo, Interference competition among burying beetles (Silphidae, Nicrophorus). Ecol. Entomol. 15, 347–355 (1990).

34. S. T. Trumbo, Infanticide, sexual selection and task specialization in a biparental burying beetle. Anim. Behav. 72, 1159–1167 (2006).

35. B. Brodie, R. Gries, A. Martins, S. VanLaerhoven, G. Gries, Bimodal cue complex signifies suitable oviposition sites to gravid females of the common green bottle fly. Entomol. Exp. Appl. 153, 114–127 (2014).

36. F. Verheggen, et al., The Odor of Death: An Overview of Current Knowledge on Characterization and Applications. BioScience 67, 600–613 (2017).

37. C. Von Hoermann, et al., Linking bacteria, volatiles and insects on carrion: the role of temporal and spatial factors regulating inter-kingdom communication via volatiles. R. Soc. Open Sci. 9, 220555 (2022).

38. G. Yan, et al., Behavior and Electrophysiological Response of Gravid and Non-Gravid Lucilia cuprina (Diptera: Calliphoridae) to Carrion-Associated Compounds. J. Econ. Entomol. 111, 1958–1965 (2018).

39. M. Kaltenpoth, S. Steiger, Unearthing carrion beetles’ microbiome: characterization of bacterial and fungal hindgut communities across the S ilphidae. Mol. Ecol. 23, 1251–1267 (2014).

40. P. Heise, et al., Antibiotic-Producing Beneficial Bacteria in the Gut of the Burying Beetle Nicrophorus vespilloides. Front. Microbiol. 10, 1178 (2019).

41. K. Brinkrolf, et al., Genomic analysis of novel Yarrowia-like yeast symbionts associated with the carrion-feeding burying beetle Nicrophorus vespilloides. BMC Genomics 22, 323 (2021).

42. N. J. Hayward, T. H. Jeavons, A. J. Nicholson, A. G. Thornton, Methyl mercaptan and dimethyl disulfide production from methionine by Proteus species detected by head-space gas-liquid chromatography. J. Clin. Microbiol. 6, 187–194 (1977).

43. K. E. Seefeldt, B. C. Weimer, Diversity of Sulfur Compound Production in Lactic Acid Bacteria. J. Dairy Sci. 83, 2740–2746 (2000).

44. M. R. H. Hurst, C. Van Koten, T. A. Jackson, Pathology of Yersinia entomophaga MH96 towards Costelytra zealandica (Coleoptera; Scarabaeidae) larvae. J. Invertebr. Pathol. 115, 102–107 (2014).

45. G. Zhu, et al., Identification and Pathogenicity of a New Entomopathogenic Fungus, Mucor hiemalis (Mucorales: Mucorales), on the Root Maggot, Bradysia odoriphaga (Diptera: Sciaridae). J. Insect Sci. 22, 2 (2022).

46. S. T. Trumbo, M. Kon, D. Sikes, The reproductive biology of Ptomascopus morio, a brood parasite of Nicrophorus. J. Zool. 255, 543–560 (2001).

47. S. T. Trumbo, D. S. Sikes, Resource concealment and the evolution of parental care in burying beetles. J. Zool. 315, 175–182 (2021).

48. E. J. Watson, C. E. Carlton, Succession of Forensically Significant Carrion Beetle Larvae on Large Carcasses (Coleoptera: Silphidae). Southeast. Nat. 4, 335–346 (2005).

49. D. C. Deeming, Nest construction in mammals: a review of the patterns of construction and functional roles. Philos. Trans. R. Soc. B Biol. Sci. 378, 20220138 (2023).

50. G. J. Dury, A. P. Moczek, D. B. Schwab, Maternal and larval niche construction interact to shape development, survival, and population divergence in the dung beetle Onthophagus taurus. Evol. Dev. 22, 358–369 (2020).

51. M. M. Lambrechts, D. C. Deeming, Nest design and breeding success in intraspecific investigations of non-cavity nesting avian species. Avian Biol. Res. 18, 38–61 (2025).

52. E. Strohm, K. E. Linsenmair, Females of the European beewolf preserve their honeybee prey against competing fungi. Ecol. Entomol. 26, 198–203 (2001).

53. J. F. Odling-Smee, K. N. Laland, M. W. Feldman, Niche construction: the neglected process in evolution (Princeton University Press, 2003).

54. B. J. Callahan, et al., DADA2: High-resolution sample inference from Illumina amplicon data. Nat. Methods 13, 581–583 (2016).

55. E. Bolyen, et al., Reproducible, interactive, scalable and extensible microbiome data science using QIIME 2. Nat. Biotechnol. 37, 852–857 (2019).

56. M. Chuvochina, et al., SILVA in 2026: a global core biodata resource for rRNA within the DSMZ digital diversity. Nucleic Acids Res. 54, D334–D341 (2026).

57. K. Abarenkov, et al., The UNITE database for molecular identification and taxonomic communication of fungi and other eukaryotes: sequences, taxa and classifications reconsidered. Nucleic Acids Res. 52, D791–D797 (2024).

58. F. Hartig, DHARMa: Residual Diagnostics for Hierarchical (Multi-Level / Mixed) Regression Models. R package version 0.4.7 (2016). 10.32614/CRAN.package.DHARMa.

