## Supplementary material for "Multifaceted parental niche construction buffers microbial and competitive challenges and drives offspring dependence in burying beetles": SI Appendix

### SI Results:

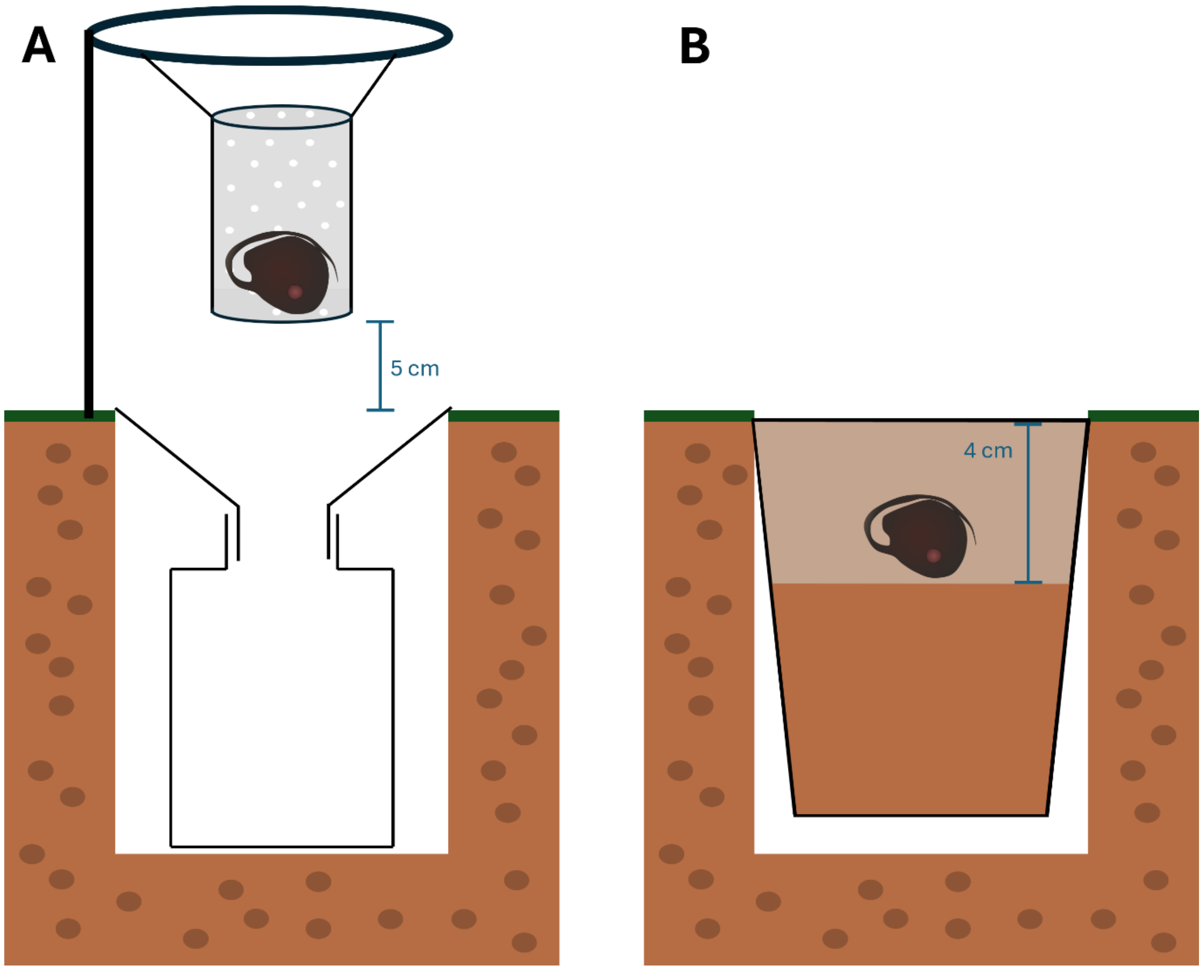


**Figure. S1: Schematic view of the trap design used in the field experiments.** A. Traps used for above-ground exposure. B. Traps used for below-ground exposure.

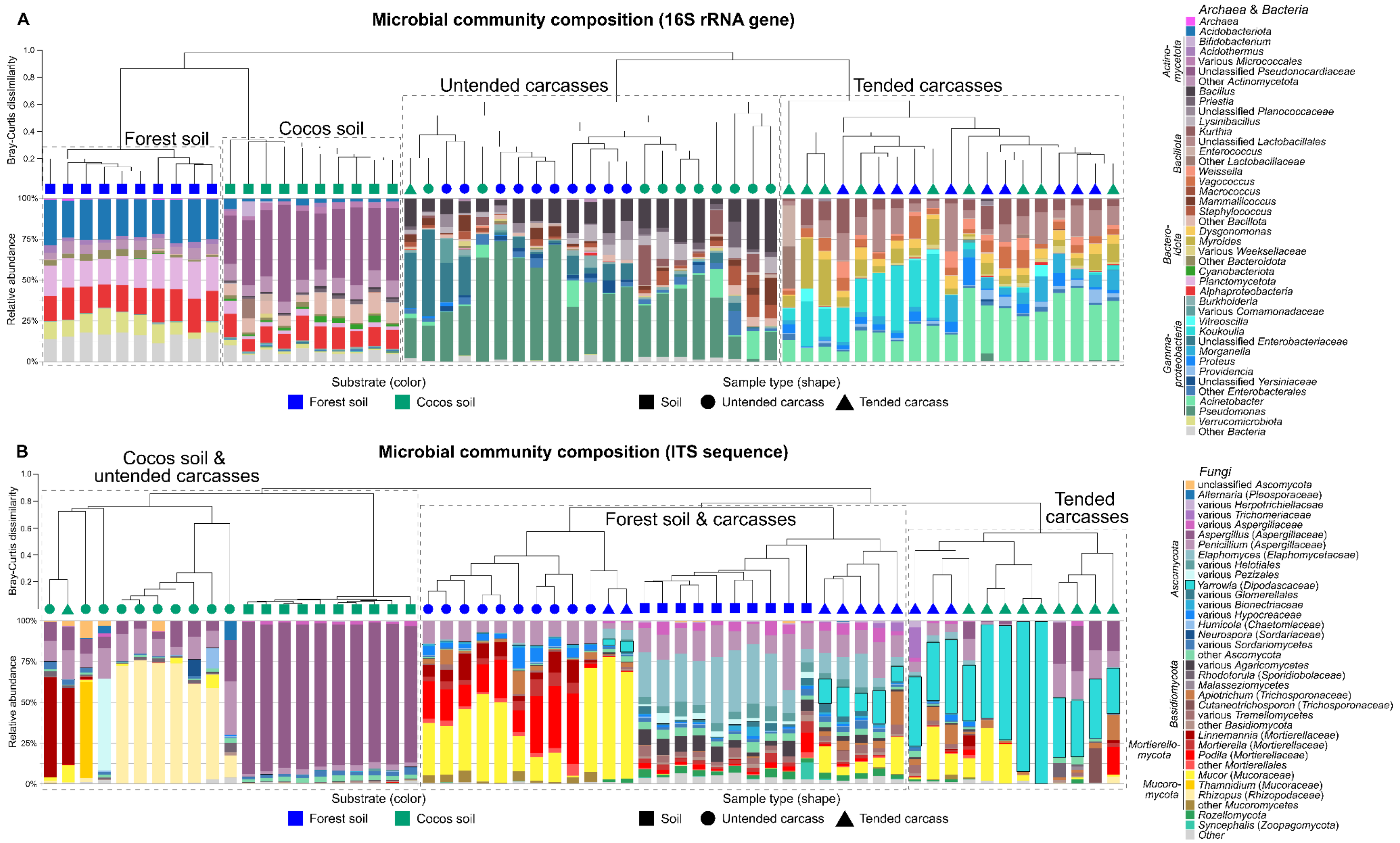


Figure S2: **Microbial community composition** based on A) 16S rRNA gene amplicons and B) fungal ITS sequences. Barplots show the relative abundances of taxonomically classified sequences. The dendrograms show the average linkage hierarchical clustering of samples based on Bray-Curtis dissimilarity of their community compositions.

**Table S1: Effects of pre-hatch care on the attraction of insect rivals.** Results of GLMMs of the effects of carcass preparation (carcass type), sampling site and day on the number of *Nicrophorus*, silphine beetles or dipteran individuals captured. Significant p-values are shown in bold.

| | | Carcass type | | | Sampling Site | | | Sampling Day | | |
|---|---|---|---|---|---|---|---|---|---|---|
| Exposure | Capture | $\chi^2$ | df | p | $\chi^2$ | df | p | $\chi^2$ | df | p |
| Above Ground | *Nicrophorus* | 0.47 | 1 | 0.49 | 0.83 | 1 | 0.36 | 1.57 | 2 | 0.46 |
| | Silphinae | 0.24 | 1 | 0.62 | 0.11 | 1 | 0.74 | 2.25 | 2 | 0.32 |
| | Diptera | 19.30 | 1 | **<0.001** | 0.03 | 1 | 0.86 | 3.57 | 2 | 0.17 |
| Below Ground | *Nicrophorus* | 12.5 | 1 | **<0.001** | 1.64 | 1 | 0.20 | 2.6 | 2 | 0.27 |

**Table S2: Effects of pre-hatch care and substrate on the composition of microbial communities of carrion.** Results of pairwise PERMANOVA comparisons (Bonferroni corrected) between microbial communities from coconut substrate and forest soil, and from tended and untended carcasses on both substrates. Significant p-values are shown in bold.

| Locus | Group 1 | Group 2 | $R^2$ | F | p |
|---|---|---|---|---|---|
| 16S rRNA | Cocos Soil | Forest Soil | 0.85 | 98.80 | **<0.001** |
| | | Untended Cocos | 0.70 | 42.57 | **<0.001** |
| | | Untended Forest | 0.80 | 70.69 | **<0.001** |
| | | Tended Cocos | 0.64 | 32.50 | **<0.001** |
| | | Tended Forest | 0.79 | 67.28 | **<0.001** |
| | Forest Soil | Untended Cocos | 0.75 | 53.80 | **<0.001** |
| | | Untended Forest | 0.83 | 88.64 | **<0.001** |
| | | Tended Cocos | 0.67 | 36.68 | **<0.001** |
| | | Tended Forest | 0.82 | 80.06 | **<0.001** |
| | Untended Cocos | Untended Forest | 0.21 | 4.86 | **<0.001** |
| | | Tended Cocos | 0.44 | 14.22 | **<0.001** |
| | | Tended Forest | 0.61 | 27.85 | **<0.001** |
| | Untended Forest | Tended Cocos | 0.57 | 23.75 | **<0.001** |
| | | Tended Forest | 0.73 | 49.20 | **<0.001** |
| | Tended Cocos | Tended Forest | 0.06 | 1.21 | 0.24 |
| ITS | Cocos Soil | Forest Soil | 0.89 | 150.16 | **<0.001** |
| | | Untended Cocos | 0.62 | 29.24 | **<0.001** |
| | | Untended Forest | 0.87 | 122.94 | **<0.001** |
| | | Tended Cocos | 0.66 | 35.97 | **<0.001** |
| | | Tended Forest | 0.76 | 56.77 | **<0.001** |
| | Forest Soil | Untended Cocos | 0.55 | 21.68 | **<0.001** |
| | | Untended Forest | 0.69 | 40.81 | **<0.001** |
| | | Tended Cocos | 0.60 | 27.15 | **<0.001** |
| | | Tended Forest | 0.45 | 14.96 | **<0.001** |
| | Untended Cocos | Untended Forest | 0.53 | 20.12 | **<0.001** |
| | | Tended Cocos | 0.38 | 11.06 | **<0.001** |
| | | Tended Forest | 0.46 | 15.27 | **<0.001** |
| | Untended Forest | Tended Cocos | 0.51 | 19.07 | **<0.001** |
| | | Tended Forest | 0.44 | 14.40 | **<0.001** |
| | Tended Cocos | Tended Forest | 0.27 | 6.71 | **<0.001** |

**Table S3: Effects of pre-hatch care and substrate on larval fitness traits.** Results of estimated marginal means (EMMeans) pairwise comparisons of larval survival and mass between tended and untended carcasses on forest soil and coconut substrate or untreated and sterilized forest soil. Significant p-values are shown in bold.

| **Larval survival – Cocos vs forest soil** | | | | | | |
|---|---|---|---|---|---|---|
| Substrate | Comparison | Odds ratio | SE | df | z.ratio | p |
| Cocos substrate | Untended vs Tended | 1.44 | 0.87 | inf | 0.61 | 0.54 |
| Forest soil | Untended vs Tended | 0.027 | 0.036 | inf | -2.73 | **0.006** |
| **Larval survival – untreated vs sterilized forest soil** | | | | | | |
| Sterilized forest soil | Untended vs Tended | 1.07 | 0.64 | inf | 0.12 | 0.91 |
| Untreated forest soil | Untended vs Tended | 0.082 | 0.067 | inf | -3.01 | **0.002** |

**Table S4: Effects of pre-hatch care on larval fitness traits of *P. morio* and *N. vespilloides*.** Results of GLMs of the effects of carcass preparation, species and their interaction of larval survival and mass after 48 hours, and at dispersal. Significant p-values are typed bold.

| | | Species | | | Carcass type | | | Species * Carcass type | | |
|---|---|---|---|---|---|---|---|---|---|---|
| Endpoint | Time | $\chi^2$ | df | p | $\chi^2$ | df | p | $\chi^2$ | df | p |
| Survival | 48 hours | 288.31 | 1 | **<0.001** | 11.17 | 1 | **<0.001** | 3.14 | 1 | 0.08 |
| | Dispersal | 299.71 | 1 | **<0.001** | 11.91 | 1 | **<0.001** | 4.17 | 1 | **0.041** |
| Mass | 48 hours | 102.49 | 1 | **<0.001** | 5.27 | 1 | **0.022** | 1.45 | 1 | 0.23 |
| | Dispersal | 151.41 | 1 | **<0.001** | 13.75 | 1 | **<0.001** | 0.05 | 1 | 0.81 |

**Table S5: Effects of pre-hatch care on larval fitness traits of *P. morio* and *N. vespilloides*.** Results of EMMeans pairwise comparisons of survival and mass of *P. morio* and *N. vespilloides* larvae between tended and untended carcasses. Significant p-values are typed bold.

| **Larval survival** | | | | | | | |
|---|---|---|---|---|---|---|---|
| Species | Time | Comparison | Odds ratio | SE | df | z.ratio | p |
| *N. vespilloides* | 48 hours | Untended vs Tended | 0.27 | 0.1 | inf | -3.48 | **<0.001** |
| *N. vespilloides* | Dispersal | Untended vs Tended | 0.21 | 0.09 | inf | -3.47 | **<0.001** |
| *P. morio* | 48 hours | Untended vs Tended | 0.78 | 0.37 | inf | -0.53 | 0.6 |
| *P. morio* | Dispersal | Untended vs Tended | 0.69 | 0.27 | inf | -0.97 | 0.33 |
| **Larval mass** | | | | | | | |
| Species | Time | Comparison | estimate | SE | df | t.ratio | p |
| *N. vespilloides* | 48 hours | Untended vs Tended | -3.44 | 1.76 | 80 | -1.96 | 0.054 |
| *N. vespilloides* | Dispersal | Untended vs Tended | -18.9 | 8.80 | 72 | -2.15 | **0.035** |
| *P. morio* | 48 hours | Untended vs Tended | -1.17 | 0.69 | 80 | -1.7 | 0.09 |
| *P. morio* | Dispersal | Untended vs Tended | -16.5 | 5.44 | 72 | -3.03 | **0.003** |

#### Table S6: Cycling conditions of the 16S rRNA and ITS2 pcr.

| Locus | Step | Duration | Temperature |
|---|---|---|---|
| 16S rRNA | Start | 3 min | 95°C |
| | Denaturing | 30 sec | 95°C |
| | Annealing | 30 sec | 55°C |
| | Extension | 30 sec | 72°C |
| | | | 25 cycles |
| | Extension | 5 min | 72°C |
| | End | | 14°C |
| ITS2 | Start | 3 min | 95°C |
| | Denaturing | 30 sec | 95°C |
| | Annealing | 30 sec | 55°C |
| | Extension | 30 sec | 72°C |
| | | | 30 cycles |
| | Extension | 5 min | 72°C |
| | End | | 14°C |

### SI Methods

#### Experimental Beetles

All beetles derived from outbred populations reared at the University of Bayreuth, Germany. *N. vespilloides* beetles originate from wild caught beetles captured in a forest near Bayreuth, Germany in 2023, *Ptomascopus morio* derived from Chiba, Japan, collected in 2023. Beetles of both species were kept in plastic containers (10 x 10 x 6 cm) filled with coconut substrate (Tropic Shop, Nordhorn, Germany) at 20°C under a 16L:8D regime. Beetles were fed with *Lucilia sericata* larvae twice a week.

#### Experimental carcasses and larvae

We established two carcass types: tended carcasses, processed by a pair of burying beetles, and untended carcasses, incubated in soil to mimic the decay status of an untended carcass at larval emergence. Carcasses were placed on either coconut substrate (Tropic Shop, Nordhorn, Germany), commonly used in our lab, or the humus-rich upper layer of forest soil collected in a mixed forest near Bayreuth, Germany.

For tended carcasses, we placed beetle pairs in plastic containers (10 × 10 × 6 cm) half-filled with the respective substrate and provided a freshly thawed mouse carcass (*Mus musculus*; 10 ± 2 g; Frostfutter.de, Krefeld, Germany). The pairs were kept at 20°C in complete darkness. After oviposition, we transferred parents and carcasses into a new container with the same substrate to separate parents and carcasses from the eggs and developing larvae. We monitored egg boxes every two hours during larval hatching (72 h post-setup) and pooled emerging larvae by substrate type in Petri dishes lined with moist filter paper. We only used tended carcasses from families with hatched larvae to ensure that pre-hatch care had been completed. We generated untended carcasses by burying freshly thawed carcasses in the respective substrate. To maintain consistency between treatments, untended carcasses were also kept at 20°C and complete darkness, transferred to new containers after the oviposition period and used in experiments as soon as larvae began to hatch.

#### Cadaveric Volatiles

We assessed how pre-hatch care by *Nicrophorus vespilloides* influences carcass VOC emissions using headspace analysis of tended (N = 24) and untended (N = 25) carcasses generated on forest soil. We collected the VOCs using a dynamic headspace method. We placed individual carcasses in commercial oven bags and pumped air through the bag at a flow rate of 200 ml / min using membrane pumps (DC12/16FK, Fürgut, Aichstetten, Germany) for 60 minutes at 22 °C. The effluent air stream passed an activated charcoal filter (Orbo 32, Supelco, Bellefonte, PA-USA) in which the volatiles emitted from the carrion source were collected. For chemical analysis, we extracted the activated charcoal from the filters and washed it in 500 µl dichloromethane (Rotisolf ≥ 99,9%, Roth, Karlsruhe, Germany) containing 20 ng/µl ethyl undecanoate (CAS-NR: 1731-86-8, Sigma-Aldrich, Taufkirchen, Germany) acting as an internal standard. We then auto-injected 1 µl per sample splitless into a GC-MS

(Shimadzu GC2030 gas chromatograph connected to Shimadzu QP2020NX mass spectrometer, Shimadzu, Duisburg, Germany). The GC contained a polar column with a length of 30 m, an inner diameter of 0.25 mm and a film thickness of 0.025 µm (RT-WAX, Restek, Bad Homburg, Germany). The temperature program of the GC oven started at 40°C, which was held for 5 min and then increased by 5°C / min up to 180°C. Helium acted as carrier gas at a linear velocity of 50 ml / min. Compounds were identified by comparison with mass spectral libraries and retention times of reference standards.

#### Field Study

We tested the effects of pre-hatch care on the attraction of insect competitors to carcasses in two field experiments conducted in May and August 2024 in a mixed forest near Betzenstein, Germany (49°55'13.9"N 11°34'20.7"E). We focused on burying beetles, other carrion beetles (Silphinae), and dipteran insects as the main competitors for carrion resources. Because burying beetles naturally bury carcasses, and burial may influence insect attraction, we exposed carcasses either above or below ground. During above-ground exposure, we placed tended and untended carcasses (each treatment N = 27) in plastic cups, mounted on plant trellises approximately 5 cm above a pitfall trap (Fig. S1 A). Perforations (1–3 mm) in the cups allowed volatile release and entry of dipteran insects into the cups, where they were trapped, enabling assessment of their attraction to the carcasses. We established 18 traps at two different sites ("Skilift": 10 traps, "Eibgrad": 8 traps) equidistant (20m) from each other. We exposed the carcasses in the field on three consecutive days in May at 12:00 and collected them 24 hours later. We identified and counted all carrion-feeding insects of the subfamily Silphinae captured in the pitfall traps. The cups containing carcasses were then frozen to allow quantification of fly visitation.

In the below-ground exposure experiment, we aimed to simulate natural burial conditions. To do so, we cut openings into the dorsal side of carcasses at hip and shoulder level prior to preparation or incubation (N = 27 per treatment), mimicking feeding holes created by parents during carcass burial (see also (1)). Carcasses were then placed in soil-filled flower cups (diameter 8.8 cm, height 10 cm) containing 6 cm of site soil and covered with an additional 4 cm of soil (Fig. S1 B). The cups were inserted into the ground so that their rims were level with the surface. We established a total of 18 soil traps at the same locations used in the previous field experiment. Carcasses were exposed in the field on three consecutive days in August, starting at 17:00, and collected at 09:00 the following morning. After collection, we identified and counted all carrion beetles present in the soil within each cup.

Across both experiments, treatments were rotated daily to avoid location-based biases, and all containers were removed and cleaned before each new trial to eliminate residual cues. Carcasses were only exposed on days with average daytime temperatures above 13 °C and without precipitation.

#### Characterization of the Carcass Microbiome

We assessed the effects of soil type and pre-hatch care on carcass-associated microbial communities using 16S rRNA and ITS amplicon sequencing of tended and untended carcasses exposed to forest soil or coconut substrate, and of the corresponding forest soil and coconut substrate samples (N = 10 per treatment and substrate). We sampled carcass-associated microbial communities after 72 h of pre-hatch care or incubation by swabbing the feeding cavities of tended carcasses, where larvae typically aggregate, and the corresponding skin region of untended carcasses to ensure comparability. We transferred the swabs immediately after sampling in RNAlater stabilizing and protection medium with immediate RNase inactivation (SigmaAldrich, Steinheim, Germany). We sampled both substrates on the day of setup (72 hours prior to carcass sampling) by transferring soil into sterile 1.5 ml reaction tube using a teaspoon. We disinfected the spoon in 70% EtOH between sampling. The soil and carcass samples were stored at -21°C until DNA extraction.

DNA was isolated using the DNAeasy Power Soil Kit (Qiagen, Hilden, Germany) according to manufacturer's guideline and stored at -21°C until use. We confirmed the presence of genomic DNA by agarose gel electrophoresis. We assessed the microbial community composition by high throughput sequencing of 16S rRNA gene fragments (V4 region) and the fungal internal transcribed spacer (ITS2) region. 16S rRNA amplicons were generated using the primers 515F-Y (2) and 806RB (3), extended at the 5'-end with Illumina overhang adapters (LIT). In particular, the 515F-Y and gITS7 primers were prepared as a phased-primer mixture where zero, one (C), or two (TC) bases were added in between the Illumina adapter and the primer sequences, respectively, and mixed in equimolar amounts. PCR was conducted with 25 cycles of amplification. Fungal ITS2 amplicons were generated using the primers *gITS7* and *ITS4* (4) with 30 amplification cycles. Detailed cycling conditions can be found in SI Tab. S7. PCR was conducted using KAPA Hifi Hot Start Ready Mix (Roche, Basel, Switzerland) according to manufacturer's guidelines and 3 µl DNA extract (5 ng/µl) as template (see Tab. S6 for cycling conditions).

Amplicons were purified using the NucleoMag NGS Clean-up and Size Selection kit (Macherey-Nagel, www.mn-net.com) as recommended by the manufacturer. Sample-specific DNA indices were added to each amplicon sample (2.5 µl of purified amplicon PCR product) in a second PCR with unique index primer combinations taken from the Nextera XT Index kit (Illumina, www.illumina.com) and quality-checked by capillary electrophoresis (Fragment Analyzer 5200, Agilent, www.agilent.com). Sample-specific libraries were combined at equimolar amounts to 16S and ITS library pools and size-selected on Pippin Prep (SAGE Science, www.sagescience.com) at a target range of 440 bp (16S library pool), or at a range from 300 to 920 bp (ITS library pool), respectively. Both library pools were combined at equimolar amounts and sequenced on an iSeq-100 device (Illumina, www.illumina.com) in 300 bp single-end mode. Raw sequence data were demultiplexed in the Illumina iSeq-100 device and stored in fastq format.

Metabarcoding bioinformatics were performed using QIIME2 (ver. amplicon_2024.10; (5)). Amplicon primers were detected and removed from the reads with Cutadapt (6) (via *q2-cutadapt*) in two consecutive steps: First, forward primer sequences (151f and gITS7) were searched and trimmed from 16S and ITS2 amplicon reads, respectively, and all untrimmed reads were discarded. Second, reverse primer sequences (806r and ITS4) were searched and trimmed from 16S and ITS2 amplicon reads, respectively, but only untrimmed reads of the 16S amplicon reads generally discarded, while untrimmed reads of the ITS2 amplicon reads were kept in the dataset due to potential longer ITS2 amplicon sequences exceeding the single-end reads length of 300 bp. The trimmed reads were denoised and filtered for chimeras with DADA2 (7) (via q2-dada2 *trim-single*) resulting in sequence features (amplicon sequencing variants, ASVs) and corresponding feature abundance tables. Potential contaminating features were identified and removed from the obtained ASV features by the prevalence method of decontam (8)(via q2-quality-control *decontam-identify*) using control samples (parallel extraction samples without added sample material, and no-template PCR samples, respectively). Taxonomic classification of ASVs were obtained using naïve-bayes trained classifiers based on the SILVA 138.2 (9) (for 16S amplicons); and UNITE 10 (10) (for ITS2 amplicons), respectively.

#### Soil sterilization

Sterilized soil was generated by autoclaving 350 ± 50 g of forest soil in autoclave bags sealed with indicator tape at 121 °C for 20 min, followed by a 10 min drying phase at 100 kPa. All bags were weighed before and after sterilization, and any water loss was compensated by adding sterile deionized water to restore initial mass. Microbial load of the substrates was assessed by plating soil suspensions on tryptic soy agar (TSA; Carl Roth, Karlsruhe, Germany). For each substrate, 10 samples were suspended in sterile deionized water (100 mg/ml) and homogenized. Samples were subsequently diluted to 1:100. Aliquots of 100 µl were spread onto TSA plates and incubated at 25 °C. Colony-forming units (CFUs) were quantified after 48 h of incubation. Soil sterilization drastically reduced the number of colony-forming units (CFUs) per gram (Wilcoxon test, W = 0, $p < 0.001$), decreasing from $3.68 \times 10^5 \pm 4.59 \times 10^5$ CFU $g^{-1}$ in forest soil to 0.00 ± 0.00 CFU $g^{-1}$ in sterilized soil.

#### Statistical analysis

All statistical analyses were performed in R (version 4.4.2). We analyzed larval survival across experiments using generalized linear models (GLMs) following a quasibinomial error distributrion. The response variable was the number of surviving larvae out of the initial brood size, modeled using *cbind(survived, not survived)* to reflect binomial sampling. In Experiment 1, the fixed effect was carcass type (tended or untended). In the experiments comparing larval performance between substrates or between species, we fitted 2 × 2 factorial models including carcass type and substrate (substrate comparison) or species (species comparison), as well as their interaction. We analyzed mean larval mass using GLMs following a gaussian error distribution, using the same predictors listed above. In cases where residual variance differed between groups, we included dispersion models (*dispformula,* within *glmmTMB* (11)) to allow the variance to depend on substrate (substrate comparison) or species (species comparison).

To determine if tended or untended carcasses can be separated based on their volatile profiles, we performed a PERMANOVA (permutational analysis of variance, *adonis2()* command in the *vegan* package (12)), based on bray curtis dissimilarities (*vegdist* and *adonis2(),* within *vegan*). We visualized differences in the profiles using NMDS plots based on bray-curtis dissimilarities (*vegdist* and *metaMDS* within *vegan*). Ordinations were performed in two dimensions, and stress values were inspected to assess ordination quality. We used Wilcoxon rank sum tests to test which substance emissions differed between carcass types.

We analyzed the effects of carcass preparation on the attraction of insect rivals using generalized linear mixed models (GLMMs(11)) with a negative binomial error distribution. The number of attracted rivals was included as the response variable, with carcass type, sampling site, and day as fixed effects. Trap ID was included as a random intercept to account for non-independence among measurements from the same trap location.

To compare the microbial community of tended and untended carcasses on forest soil and coconut substrate, we performed pairwise PERMANOVAs (pairwise.*adonis2()* within *pairwiseAdonis* (13)), based on bray curtis dissimilarities (*distance* within *phyloseq* (14)). To account for the 2 x 2 factorial design within the carcass samples, we reanalyzed the carcass communities using PERMANOVAs based on Bray Curtis dissimilarities using substrate, carcass type and their interactions as predictors. We visualized community differences among tended and untended carcasses on both substrates using NMDS plots based on Bray-Curtis dissimilarities (*distance* within *phyloseq* and *metaMDS* within *vegan*). Ordinations were performed in two dimensions, and stress values were inspected to assess ordination quality. Hierarchical clustering of samples based on microbial community composition was calculated using the *hclust* function and average linkage method based on a Bray-Curtis dissimilarity matrix (*vegdist* function within *vegan*). Ordination of samples based on their community composition was calculated using the *metaMDS* function based on Bray-Curtis dissimilarity matrix and using the *monoMDS* method with 1000 iterations.

Differential abundance of microbial taxa between different treatments were determined using *DeSeq2* (15). A taxon was considered as significantly differentially abundant between two treatments if the false-positive adjusted p-value of the Wald test was ≤ 0.05 and the indicated effect size (Log2-fold change of corrected average abundance) was bigger than 1 or smaller than -1. C. For bacteria and archaea (16S rRNA gene data), comparisons were made between all tended and all untended carcasses, since these were the two separate groups observed in hierarchical clustering with no clear separation by substrate within them (Fig. S2). For fungi (ITS data), the comparisons between tended and untended caracsses were perfomed separately for forest soil and coconut substrate, owing to the fact of primary clustering of the communities by substrate. and specifically between untended carcasses on coconut substrate vs untended carcasses on forest soil. Heatmaps of differentially abundant taxa were plotted using the *heatmap.2* function from the R package gplots (16).

All further plots were generated using the *ggplot2* (17). Model fits were validated using *DHARMa* (18), no major violations were detected.

#### References for Supporting Information


1. S. T. Trumbo, P. K. B. Philbrick, J. Stökl, S. Steiger, Burying Beetle Parents Adaptively Manipulate Information Broadcast from a Microbial Community. *Am. Nat.* **197**, 366–378 (2021).

2. A. E. Parada, D. M. Needham, J. A. Fuhrman, Every base matters: assessing small subunit RRNA primers for marine microbiomes with mock communities, time series and global field samples. *Environ. Microbiol.* **18**, 1403–1414 (2016).

3. J. G. Caporaso, *et al.*, Ultra-high-throughput microbial community analysis on the Illumina HiSeq and MiSeq platforms. *ISME J.* **6**, 1621–1624 (2012).

4. K. Ihrmark, *et al.*, New primers to amplify the fungal ITS2 region - evaluation by 454-sequencing of artificial and natural communities. *FEMS Microbiol. Ecol.* **82**, 666–677 (2012).

5. E. Bolyen, *et al.*, Reproducible, interactive, scalable and extensible microbiome data science using QIIME 2. *Nat. Biotechnol.* **37**, 852–857 (2019).

6. M. Martin, Cutadapt removes adapter sequences from high-throughput sequencing reads. *EMBnet.journal* **17**, 10 (2011).

7. B. J. Callahan, *et al.*, DADA2: High-resolution sample inference from Illumina amplicon data. *Nat. Methods* **13**, 581–583 (2016).

8. N. M. Davis, D. M. Proctor, S. P. Holmes, D. A. Relman, B. J. Callahan, Simple statistical identification and removal of contaminant sequences in marker-gene and metagenomics data. *Microbiome* **6**, 226 (2018).

9. M. Chuvochina, *et al.*, SILVA in 2026: a global core biodata resource for rRNA within the DSMZ digital diversity. *Nucleic Acids Res.* **54**, D334–D341 (2026).

10. K. Abarenkov, *et al.*, The UNITE database for molecular identification and taxonomic communication of fungi and other eukaryotes: sequences, taxa and classifications reconsidered. *Nucleic Acids Res.* **52**, D791–D797 (2024).

11. M. McGillycuddy, D. I. Warton, G. Popovic, B. M. Bolker, Parsimoniously Fitting Large Multivariate Random Effects in glmmTMB. *J. Stat. Softw.* **112** (2025).

12. J. Oksanen, *et al.*, vegan: Community Ecology Package. https://doi.org/10.32614/CRAN.package.vegan. Deposited 2024.

13. M. Martinez Arbizu, pairwiseAdonis: Pairwise Multilevel Comparison using Adonis. (2017).

14. P. J. McMurdie, S. Holmes, phyloseq: An R package for reproducible interactive analysis and graphics of microbiome census data. *PLOS ONE* **8**, e61217 (2013).

15. M. I. Love, W. Huber, S. Anders, Moderated estimation of fold change and dispersion for RNA-seq data with DESeq2. *Genome Biol.* **15**, 550 (2014).

16. G. R. Warnes, *et al.*, gplots: Various R Programming Tools for Plotting Data. https://doi.org/10.32614/CRAN.package.gplots. Deposited 30 May 2005.

17. H. Wickham, “Data Analysis” in *Ggplot2*, Use R!, (Springer International Publishing, 2016), pp. 189–201.

18. F. Hartig, DHARMa: Residual Diagnostics for Hierarchical (Multi-Level / Mixed) Regression Models. R package version 0.4.7 (2016). https://doi.org/10.32614/CRAN.package.DHARMa.
